# A rapidly adaptable biomaterial vaccine for SARS-CoV-2

**DOI:** 10.1101/2020.07.07.192203

**Authors:** Fernanda Langellotto, Benjamin T. Seiler, Jingyou Yu, Mark J. Cartwright, Des White, Chyenne Yeager, Michael Super, Edward J. Doherty, Dan H. Barouch, David J. Mooney

## Abstract

The global COVID-19 pandemic motivates accelerated research to develop safe and efficacious vaccines. To address this need, we leveraged a biomaterial vaccine technology that consists of mesoporous silica rods (MSRs) that provide a sustained release of granulocyte-macrophage colony-stimulating factor (GM-CSF) and adjuvants to concentrate and mature antigen-presenting cells at the vaccine site. Here we explored the humoral responses resulting from the use of monophosphoryl lipid A (MPLA) as the adjuvant and SARS-CoV-2 spike proteins S1, S2, the nucleocapsid (N) protein, and receptor binding domain (RBD) as the target antigens. The dose of antigen and impact of pre-manufacturing of vaccines as versus loading antigen just-in-time was explored in these studies. Single shot MSR vaccines induced rapid and robust antibody titers to the presented antigens, even without the use of a boost, and sera from vaccinated animals demonstrated neutralizing activity against a SARS-CoV-2 pseudovirus. Overall, these results suggest the MSR vaccine system may provide potent protective immunity when utilized to present SARS-CoV-2 antigens.

## INTRODUCTION

In late December 2019, a cluster of pneumonia cases of unknown etiology was announced in Wuhan, China (*1,2*). In January 2020, the Chinese Center for Disease Control and Prevention (CDC) identified that a novel coronavirus, known as Severe Acute Respiratory Syndrome Coronavirus 2 (SARS-CoV-2), was the causative agent of a new respiratory tract disease, now called coronavirus disease 2019 (COVID-19) (*3,4*). COVID-19 rapidly advanced from an epidemic to a pandemic, and as of July 6, 2020 the World Health Organization (WHO) has reported 11,327,790 confirmed cases, resulting in 532,340 deaths worldwide (*5*).

Many strategies to combat COVID-19 are currently being implemented; among them include contact tracing, convalescent plasma therapy, anti-viral drug therapy, and vaccine development (*6-9*). Here, we focus on vaccine development. SARS-CoV-2 contains many structural components that provide promising target candidates, including the spike (S) proteins, nucleocapsid (N) protein, and receptor binding domain (RBD) (*10,11*). The S protein is of particular interest due to its demonstrated ability to induce potent neutralizing antibodies and T-cell responses against SARS-CoV in previous studies (*12*-*14*). Two functional subunits comprise the S protein – S1, which contains the RBD that is critical for interacting with the host cell receptor, angiotensin-converting enzyme 2 (ACE2), and S2, which mediates the fusion of viral and host cell membranes to release RNA for replication (*15,16*). The sequence identity of the S protein between SARS-CoV and SARS-CoV-2 is on the order of 76%, while RBD epitope homology is around 74%, and both respective virus types utilize ACE2 for viral entry (*17*).

Here we explored the ability of a biomaterial vaccine based on mesoporous silica rods (MSRs), which upon subcutaneous injection will spontaneously assemble to form a 3D macroporous structure, to generate humoral responses against SARS-CoV-2 relevant antigens. Granulocyte-macrophage colony-stimulating factor (GM-CSF), a potent cytokine, is incorporated on the MSRs, and its sustained release leads to the accumulation of large numbers of antigen presenting cells (APCs) at the vaccine site (*18*). The MSRs are also loaded with an adjuvant, and the sustained adjuvant release leads to maturation of the accumulated APCs, and a sustained trafficking of antigen-loaded APCs to the draining lymph node where they present antigen to B and T cells (*18*). A single injection of the MSR vaccine delivering protein or peptide antigens elicits persistent germinal center B-cell activity (>3 weeks), highly elevated antibody responses (>12 months), superior memory B-cell generation, and potent CTL responses (*19,20*) as well as an adjuvant and antigen. We have previous shown the broad-spectrum capability of this vaccine technology, demonstrating immunogenicity and efficacy against viral and bacterial pathogens (*18*-*21*).

To explore the ability of the MSR vaccine to induce potent, SARS-CoV-2 relevant humoral responses in rodents, here we loaded with the SARS-CoV-2 antigens, and the adjuvant monophosphoryl lipid A (MPLA), a TLR-4 agonist (*22*). The MSR vaccine offers a number of advantages over traditional vaccines. First, it is modular, allowing for a highly adaptable, plug-and-play nature for inclusion of various antigens and adjuvants. This feature was exploited in the current study by exploring the inclusion of several combinations of different protein antigens - S1, S2, N, and RBD. This modularity also potentially allows a rapid response to pandemics, ease of manufacture and production scale up. The dose of antigen was varied in these studies, and vaccines in which the antigens were pre-loaded onto MSRs before storage and subsequent vaccination, as versus added just before vaccination, were compared to further demonstrate the versatility of this technology. Our results demonstrate the development of rapid and robust antibody titers against all three SARS-CoV-2 proteins that were tested, in all vaccine configurations. We also demonstrate the antibodies induced by our vaccines confer functional neutralization against a SARS-CoV-2 pseudovirus.

## MATERIALS AND METHODS

### Vaccine components

To generate mesoporous silica rods (MSR) (∼46 μm × 4.5 μm), 4 g of the surfactant, P123 (M_n_ ∼5,800, Sigma Aldrich, USA), was dissolved in 150 g of 1.6 M Hydrochloric acid (HCl), then stirred with 8.6 g of tetraethyl orthosilicate (98%, Sigma Aldrich, USA) at 40°C for 20 hrs, followed by aging at 100°C for 24 hrs. The surfactant was then extracted by refluxing the particles for 24 hrs in 1% HCl in 70% ethanol. The resulting MSR particles were filtered, washed with 70% ethanol, and dried. MSR morphology was measured and determined using optical microscopy and scanning electron microscopy. Pore volume, pore size, and surface area were analyzed by N_2_ adsorption/desorption isotherms. Granulocyte-macrophage colony-stimulating factor (GM-CSF) was purchased from Peprotech, USA (315-03). Monophosphoryl lipid A (MPLA) derived from *Salmonella minnesota* R595 was purchased from Invivogen, USA (vac-mpla). SARS-CoV-2 antigens included spike protein subunits, S1 (40591-V08H), S2 (40590-V08B), and nucleocapsid (N) protein (40588-V08B) was purchased from Sino Biological, USA, and the receptor binding domain (RBD) protein (NR-52306) was purchased from BEI resources, USA.

### Vaccine manufacture

The vaccine formulations for each group are outlined in **Table 1**. To make the complete vaccine, 5 mg of MSR was added to a vial followed by 1 μg of GM-CSF, 25 μg of MPLA and the respective SARS antigen. For Group A1, the antigens were 1 µg of S1, S2, and N; for Group B the antigen were 1 µg of S1, S2, and RBD; for Group C the antigens were 5 µg of S1, S2, and RBD; and for Group D the antigen was 1 µg of RBD. Next, 0.5 ml water for injection (WFI, HyClone, USA) was added to the vial, and the resulting slurry was vortexed and mixed overnight (∼15 hrs) using a HulaMixer (ThermoScientific, USA) set at 45 rpm. Finally, the mixture was frozen at -80°C and lyophilized for a minimum of 48 hrs **(Fig. 1A)**. To make the sham and shell vaccines, 5 mg of MSR was added to a vial followed by 1 μg of GM-CSF and 25 μg of MPLA. 0.5 ml WFI was added to the vial, and the resulting slurry was vortexed and mixed overnight (∼15 hrs) using a HulaMixer set at 45 rpm. Next, the mixture was frozen at -80°C and lyophilized for a minimum of 48 hrs. Antigens in Group A2 (1 µg of S1, S2, and N) were added to shell vaccines and allowed to mix 30 min before immunization. No antigen was added to sham vaccines **(Fig. 1B**).

**Table 1.**
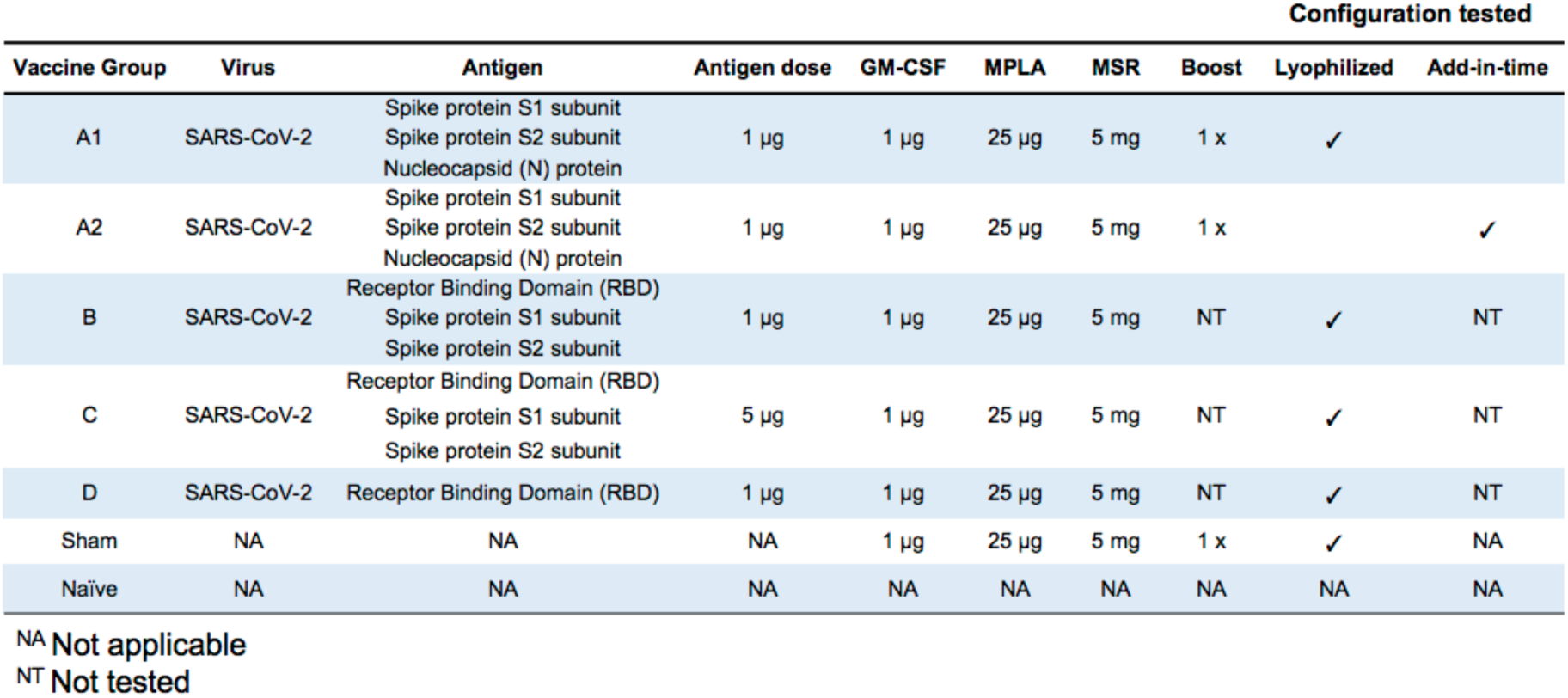
Vaccine formulations tested *in vivo*.

| Vaccine Group | Virus | Antigen | Antigen dose | GM-CSF | MPLA | MSR | Boost | Configuration tested |  |
| --- | --- | --- | --- | --- | --- | --- | --- | --- | --- |
|  |  |  |  |  |  |  |  | Lyophilized | Add-in-time |
| A1 | SARS-CoV-2 | Spike protein S1 subunit<br>Spike protein S2 subunit<br>Nucleocapsid (N) protein | 1 µg | 1 µg | 25 µg | 5 mg | 1 x | ✓ |  |
| A2 | SARS-CoV-2 | Spike protein S1 subunit<br>Spike protein S2 subunit<br>Nucleocapsid (N) protein | 1 µg | 1 µg | 25 µg | 5 mg | 1 x |  | ✓ |
| B | SARS-CoV-2 | Receptor Binding Domain (RBD)<br>Spike protein S1 subunit<br>Spike protein S2 subunit | 1 µg | 1 µg | 25 µg | 5 mg | NT | ✓ | NT |
| C | SARS-CoV-2 | Receptor Binding Domain (RBD)<br>Spike protein S1 subunit<br>Spike protein S2 subunit | 5 µg | 1 µg | 25 µg | 5 mg | NT | ✓ | NT |
| D | SARS-CoV-2 | Receptor Binding Domain (RBD) | 1 µg | 1 µg | 25 µg | 5 mg | NT | ✓ | NT |
| Sham | NA | NA | NA | 1 µg | 25 µg | 5 mg | 1 x | ✓ | NA |
| Naïve | NA | NA | NA | NA | NA | NA | NA | NA | NA |

<sup>NA</sup> Not applicable
<sup>NT</sup> Not tested

**Figure 1.**
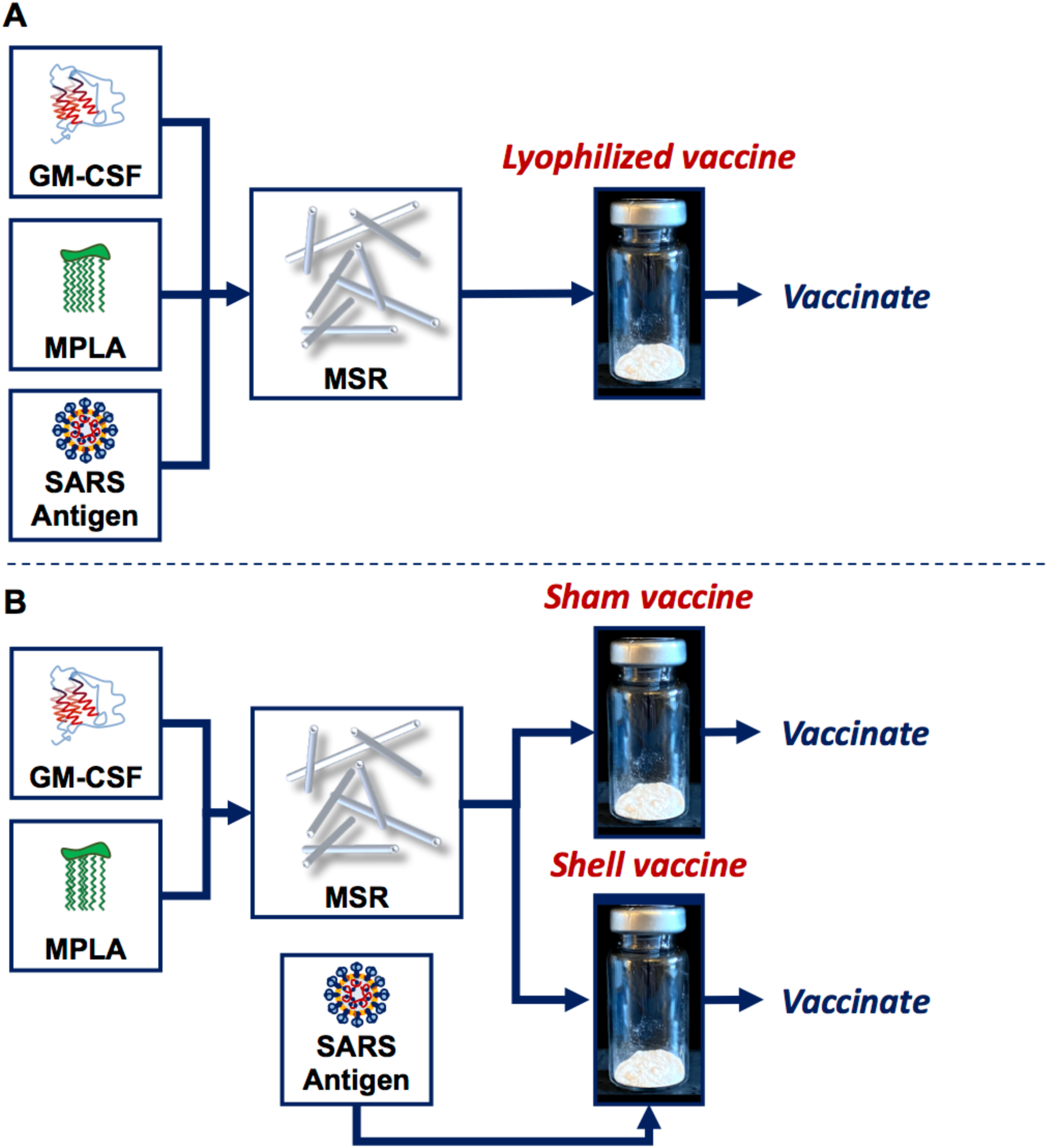
MSR vaccines can be manufactured with antigen added before lyophilization or added after, prior to vaccination. **(A)** Granulocyte-macrophage colony-stimulating factor (GM-CSF), monophosphoryl lipid A (MPLA) and the SARS antigens were adsorbed onto the biomaterial scaffold, mesoporous silica rods (MSR). The mixture was then frozen and lyophilized (lyophilized vaccine) until ready for vaccination. **(B)** GM-CSF and MPLA were adsorbed onto MSR, which was then frozen and lyophilized (sham and shell vaccine). Prior to vaccination, the SARS antigens were added just-in-time and adsorbed on shell vaccines.

### Animals and vaccination

Eight-week-old BALB/c female mice (n=10 per group) were vaccinated subcutaneously in the flank using an 18G needle. PBS was used as the unvaccinated control (naïve). All the mice were bled from submandibular vein. After collection, the whole blood was allowed to clot at room temperature for 30 min. To remove the clot, blood was centrifuged at 15,000 rpm for 15 minutes. The supernatant representing serum was aliquoted and stored at appropriate temperature. Four weeks after the primary immunization, mice were boosted subcutaneously with the same dose as the initial vaccine. Mice are maintained under specific pathogen-free conditions at Harvard University, and all experiments are conducted in accordance with animal use guidelines and protocols approved by the Harvard University Animal Care and Use Committee (IACUC).

### Antibody titers

To evaluate the IgG antibody response against SARS-CoV-2 proteins, polystyrene 96-well high-bind plates (Santa Cruz Biotechnology, USA) were coated with 100 µl of 1 µg/mL of S1, S2, and RBD in PBS. A standard curve was applied to each plate starting with 100 µl of 50 ng/ml (015-000-003, Jackson ImmunoResearch, USA) and two-fold serially diluted down, leaving the last wells as just PBS. Plates were left to incubate overnight at 4°C. The following day, the plates were washed and blocked with blocking buffer (1% BSA in TBST 5mM CaCl2) for 1 hr. Sera was diluted 1:100, 1:1,000, 1:10,000 in blocking buffer and 100 µl was added to the plates, then incubated for 1.5 hrs at 37°C on a plate shaker at 450 rpm. After sera incubation, the plates were washed and Peroxidase-AffiniPure F(ab’)2 Fragment Goat anti-mouse IgG (111-036-047, Jackson ImmunoResearch, USA) diluted to 1:20,000 in blocking buffer was added to every well and incubated for 1 hr using a plate shaker set to 450 rpm at room temperature. The plates were washed and 100 µl TMB Ultra (34028, ThermoFisher, USA) was added to all the wells and statically incubated under aluminum foil for 15 min. The reaction was quenched using 50 µl of 1M sulfuric acid and measured at OD 450nm. Samples were run in duplicates and standards in triplicate. All plate washes were done in triplicate with 200 µl of TBST 5mM CaCl_2_ using a BioTek microplate washer. Standard curve OD readouts on each plate were used to create a 4PL-sigmoidal curve to interpolate sample data, which was analyzed, processed, and presented using GraphPad Prism 8 (version 8.1.2).

### Pseudovirus neutralization assay

The SARS-CoV-2 pseudoviruses expressing a luciferase reporter gene were generated in an approach similar to as described previously (*23*). Briefly, the packaging construct psPAX2 (AIDS Resource and Reagent Program), luciferase reporter plasmid pLenti-CMV Puro-Luc (Addgene, USA), and spike protein expressing pcDNA3.1-SARS CoV-2 SΔCT were co-transfected into HEK293T cells with calcium phosphate. The supernatants containing the pseudotype viruses were collected 48 hrs post-transfection; pseudotype viruses were purified by filtration with 0.45 µm filter. To determine the neutralization activity of the antisera from vaccinated animals, HEK293T-hACE2 cells were seeded in 96-well tissue culture plates at a density of 1.75 x 10^4^ cells/well overnight. Two-fold serial dilutions of heat inactivated serum samples were prepared and mixed with 50 µL of pseudovirus. The mixture was incubated at 37°C for 1 hr before adding to HEK293T-hACE2 cells. Forty-eight hrs after infection, cells were lysed in Steady-Glo Luciferase Assay (Promega, USA) according to the manufacturer’s instructions. SARS-CoV-2 neutralization titers were defined as the sample dilution at which a 50% reduction in relative light units (RLU) was observed relative to the average of the virus control wells.

### Statistical analysis

All measured values are reported as means ± standard deviation (SD). Significant differences between groups were determined using Two-tailed Students T-test with a two-sample unequal variance (heteroscedastic) parameter.

## RESULTS

The facile manufacturing process of the MSR vaccines, and their plug-and-play nature allowed vaccines to be fabricated and mice to be vaccinated within seven days of obtaining the protein antigens.

### MSR vaccines induce SARS-CoV-2 protein specific IgG responses

The anti-S1, S2 and RBD IgG responses resulting from vaccination with Group A1 (MSR, S1/S2/N, GM-CSF, MPLA), Group B (MSR, RBD/S1/S2 1 µg, GM-CSF, MPLA), Group C (MSR, RBD/S1/S2 5 µg, GM-CSF, MPLA), and D (MSR, RBD 1 µg, GM-CSF, MPLA) were first quantified. All four of these groups were prepared by adding antigens to the MSR vaccines during fabrication, lyophilizing and storing the vaccines, and subsequently utilizing for vaccination. Negative controls in these studies included non-antigen containing shams (MSR, GM-CSF, MPLA) as well as unvaccinated naïve mice. Vaccination with Group A1 vaccines resulted in appreciable IgG levels by day 14, and these levels continued to climb until plateauing at day 28 **(Fig. 2A)**. The IgG levels remained at a similar level for the duration of the study, even though these mice received a boost at day 28. IgG levels in mice vaccinated with Groups B, C, and D vaccines increased by two orders of magnitude over the sham control by day 14 **(Fig. 2B)**, and by day 28 were greater than or equal to 1 x 10^4^ ng/ml; these levels were maintained for the duration of the study. Strikingly, these mice only received a single vaccination at day 0, with no subsequent boost. These results also demonstrated that a 5x vs 1x dose of antigen (Group C vs Group B) induced only a slightly greater IgG response **(Fig. 2B)**. The incorporation of a single antigen (RBD; Group D) led to similar anti-RBD and -SI IgG antibody responses as vaccines incorporating multiple antigens **(Fig. 2B)**. Not surprisingly, this vaccine did not induce any IgG response to S2, as the RBD is not part of the S2 subunit of the S protein.

**Figure 2.**
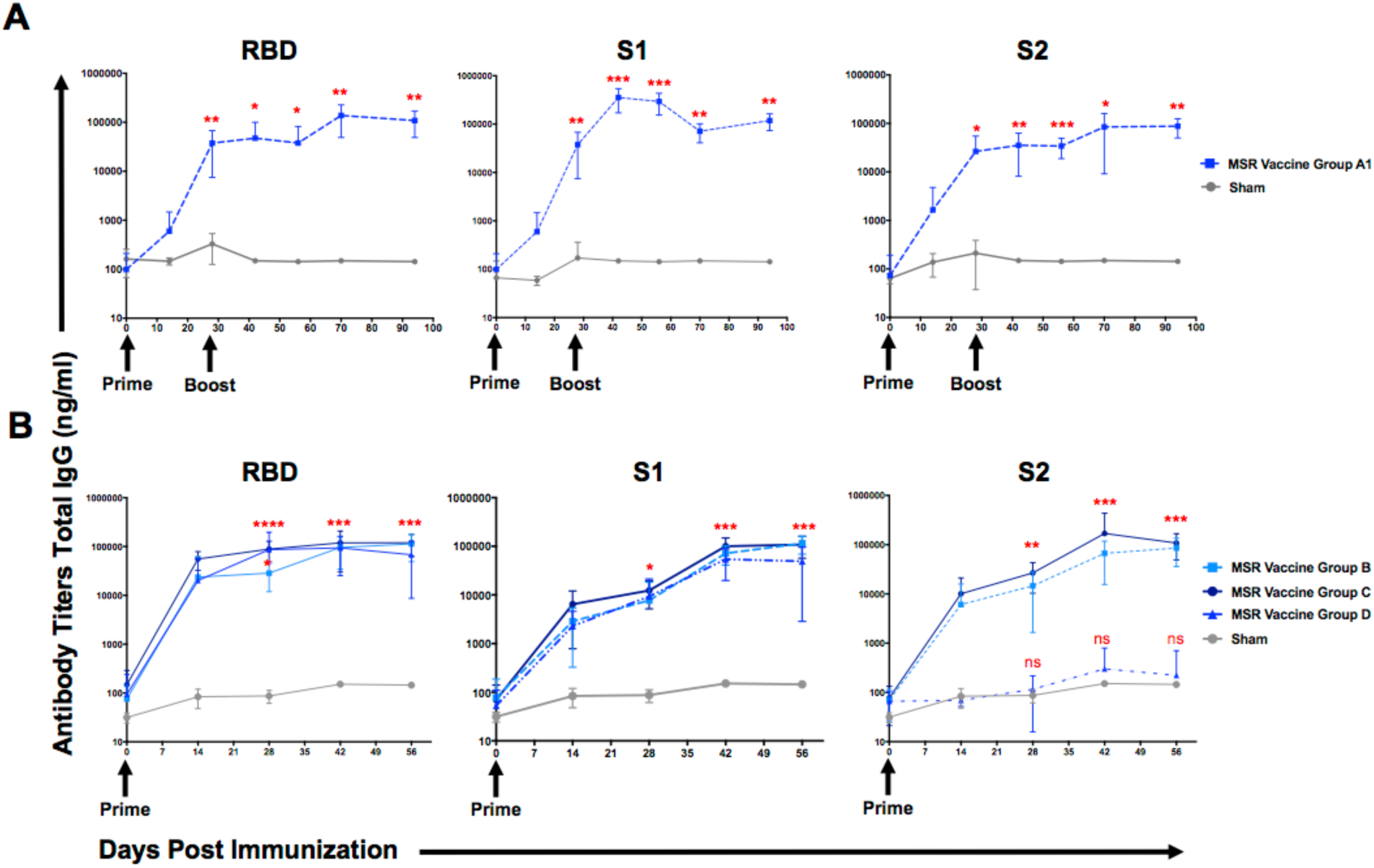
MSR lyophilized vaccines induced rapid humoral immune responses in mice. Humoral immuneresponses were assessed by lgG binding ELISA against plates coatedwith RBD, S1, and S2. **(A)** Group A1 (MSR, S1/S2/N, GM-CSF, MPLA) lyophilized and sham (MSR, GM-CSF, MPLA) vaccines were administered at day 0 (prime) and boosted on day 28 with the same formulation (n=10 per vaccine). **(B)** Group B (MSR, RBD/S1/S2 1 µg, GM-CSF, MPLA), C (MSR, RBD/S1/S2 5 µg, GM-CSF, MPLA), and D (MSR, RBD 1 µg, GM-CSF, MPLA) lyophilized and sham vaccines were administeredat day 0 with no boost (n=10 per vaccine). Each sample time point was performed in triplicate and plotted as the mean with standard deviation. *, **, ***, and **** represents p<0.05, p<0.01, p<0.001 and p<0.0001, respectively.

In addition to a standard, lyophilized vaccine configuration, an alternative method of adding antigen to the MSR vaccine was explored. In this approach, the antigens are added to shell vaccines (MSR, GM-CSF, MPLA) 30 minutes prior to immunization **(Table 1 Group A2)**. This approach allows pre-manufacturing and storage of the vaccine, minus antigen, and antigens that are identified can be readily incorporated at time of vaccination. This vaccine configuration induced no measurable IgG response by day 14, and only approximately one order of magnitude increases over the sham control by day 28 **(Fig. 3)**. However, after boosting, this vaccine formulation induced similar levels of IgG to the lyophilized vaccines (1 x 10^4^ ng/ml) against RBD and S1, but an order of magnitude lower response against S2 **(Fig. 3)**. Similar to the lyophilized vaccine configuration, IgG levels plateaued on day 42 and remained constant for the duration of the study.

**Figure 3.**
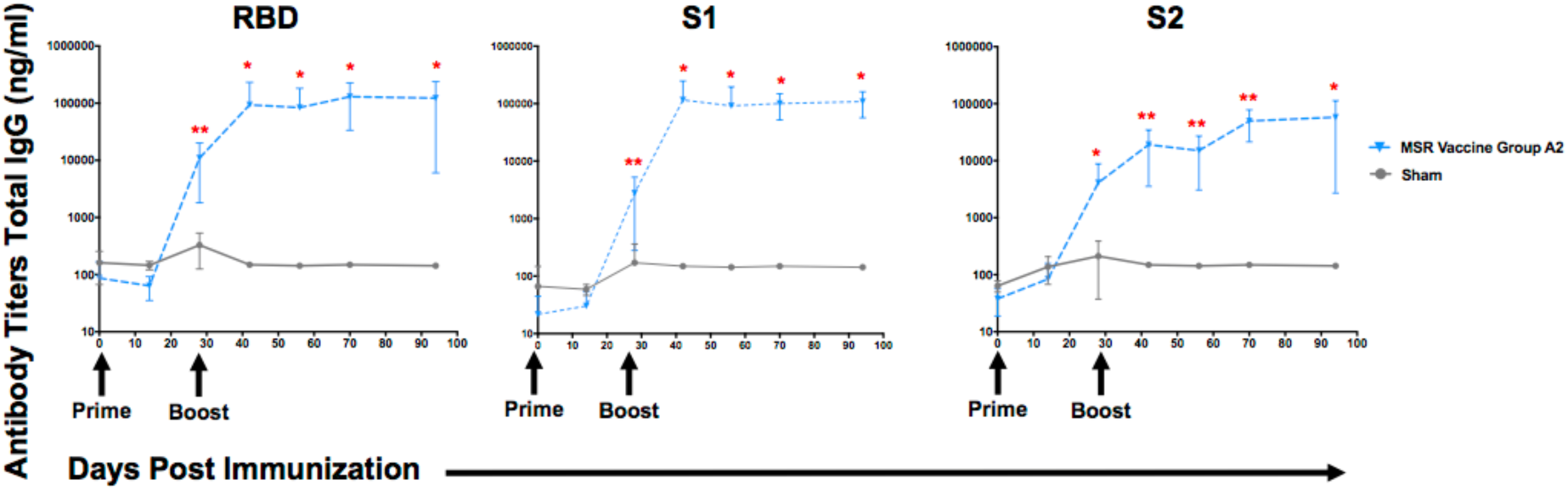
MSR add-in-time vaccines induced humoral immune responses in mice. Humoral immune responses were assessed by lgG binding ELISA against plates coated with RBD, S1, and S2. Group A2 (MSR, S1/S2/N, GM-CSF, MPLA) add-in-time and sham (MSR, GM-CSF, MPLA) vaccines were administered at day 0 (prime) and boosted on day 28 with the same formulation (n=10 per vaccine). Each sample time point was performed in triplicate and plotted as the mean with standard deviation. *, and ** represents p<0.05 and p<0.01, respectively.

### Antibodies from vaccinated mice confer neutralization of SARS-CoV-2 pseudovirus

To assess the functionality of the antibody responses induced by the MSR vaccines, sera from vaccinated mice was tested for ability to neutralize a SARS-CoV-2 pseudovirus. Sera from specific time points and vaccination conditions was analyzed **(Table 2)**. Sera from Group A1 lyophilized vaccines demonstrated minimal activity at day 28, but by day 42 and 56 the median values of antibody neutralization titers were 557 and 656, respectively **(Fig. 4A)**. By comparison, sera from Group B, C, and D vaccinated animals conferred significant neutralization at day 28 **(Fig. 4B)**. The 5x antigen dose performed slightly better, generating more neutralizing activities than the 1x dose (Group C vs Group B; median titer 445 vs 143). Sera from mice vaccinated with the RBD antigen alone (Group D) also demonstrated substantial neutralization. Group A2 add-in-time vaccines, which led to a slower induction of IgG responses than Group A1, were next analyzed. These also lead to a slower increase in neutralization activity as at day 28 the neutralization activity was less than that of the lyophilized vaccine (Group A1). However, after boosting, the neutralization titers in sera on days 42 and 56 had similar values to those from Group A1 **(Fig. 5)**. It is worth noting however, that we did see more variability with the add-in-time configuration of Group A2 vaccines, as compared to the pre-manufactured, lyophilized vaccines (Group A1).

**Table 2.** Vaccine time points selected for pseudovirus neutralization testing.

| Vaccine Group | Configuration | Virus | Antigen | Antigen dose | Pre-Boost |  | Post-Boost |  |
| --- | --- | --- | --- | --- | --- | --- | --- | --- |
|  |  |  |  |  | Day 14 | Day 28 | Day 42 | Day 56 |
| A1 | Lyophilized | SARS-CoV-2 | Spike protein S1 subunit<br>Spike protein S2 subunit<br>Nucleocapsid (N) protein | 1 µg | 10 samples | 10 samples | 10 samples | 10 samples |
| A2 | Add-in-time | SARS-CoV-2 | Spike protein S1 subunit<br>Spike protein S2 subunit<br>Nucleocapsid (N) protein | 1 µg | NT | 10 samples | 10 samples | 10 samples |
| B | Lyophilized | SARS-CoV-2 | Receptor Binding Domain (RBD)<br>Spike protein S1 subunit<br>Spike protein S2 subunit | 1 µg | NT | 10 samples | NT | NT |
| C | Lyophilized | SARS-CoV-2 | Receptor Binding Domain (RBD)<br>Spike protein S1 subunit<br>Spike protein S2 subunit | 5 µg | NT | 10 samples | NT | NT |
| D | Lyophilized | SARS-CoV-2 | Receptor Binding Domain (RBD) | 1 µg | NT | 10 samples | NT | NT |
| Sham | Lyophilized | NA | NA | NA | NT | 10 samples | NT | NT |
| Naïve | NA | NA | NA | NA | NT | 10 samples | NT | NT |

<sup>NA</sup> Not applicable
<sup>NT</sup> Not tested

**Figure 4.**
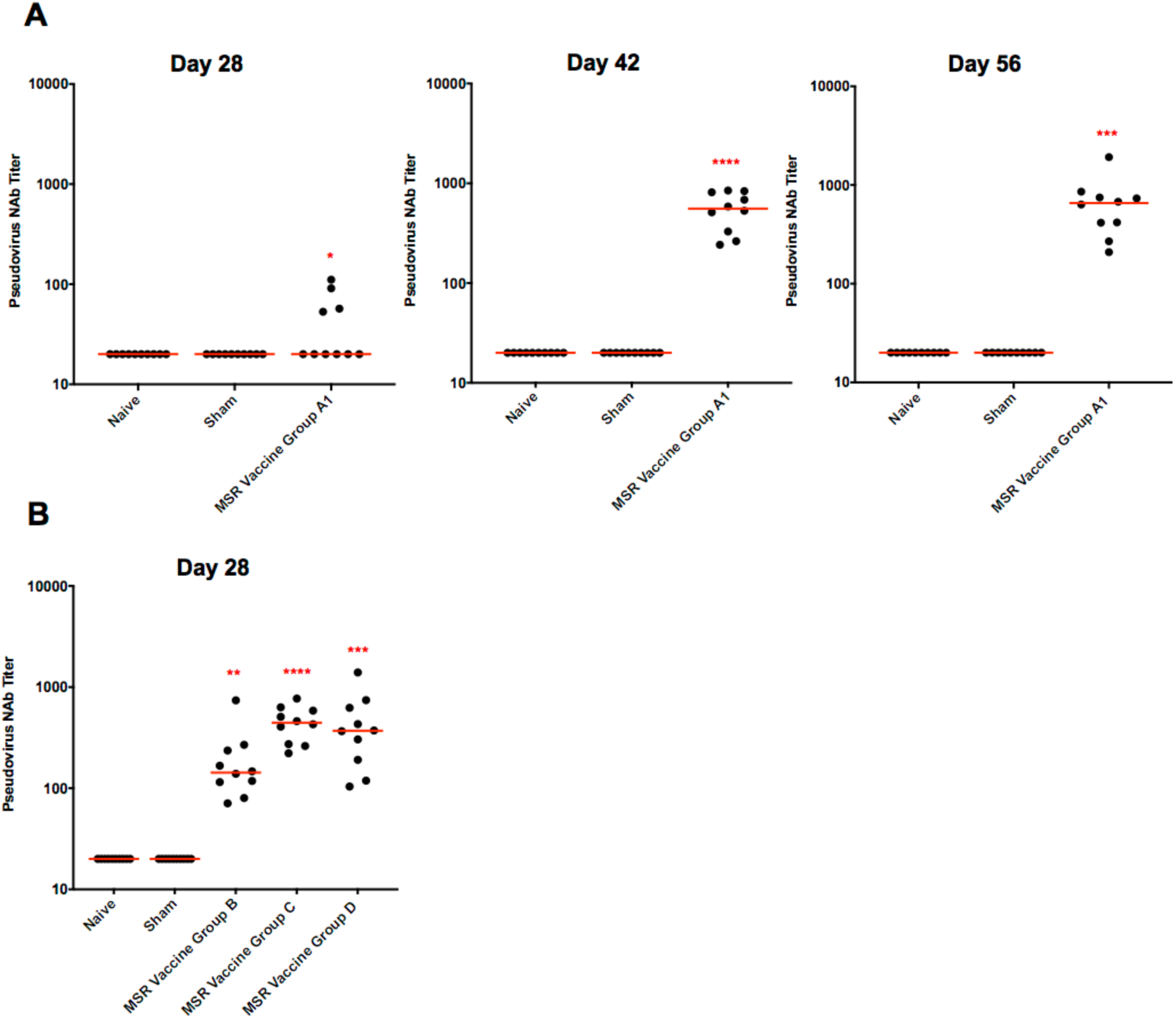
MSR lyophilized vaccines induced neutralizing antibodies (Nab) against a SARS-CoV-2 pseudovirus. Functional antibodies from sera of vaccinated mice were assessed by *in vitro* pseudovirus neutralization assay for **(A)** Group A1 (MSR, S1/S2/N, GM-CSF, MPLA) lyophilized and sham (MSR, GM-CSF, MPLA) vaccines on day 28, 42, and 56, and **(B)** Group B (MSR, RBD/S1/S2 1 µg, GM-CSF, MPLA), C (MSR, RBD/S1/S2 5 µg, GM-CSF, MPLA), and D (MSR, RBD 1 µg, GM-CSF, MPLA) lyophilized vaccines and sham vaccines on day 28. Naïve mice were unvaccinated controls. Samples were plotted as median values (n=10). *, **, ***, and **** represents p<0.05, p<0.01, p<0.001 and p<0.0001, respectively.

**Figure 5.**
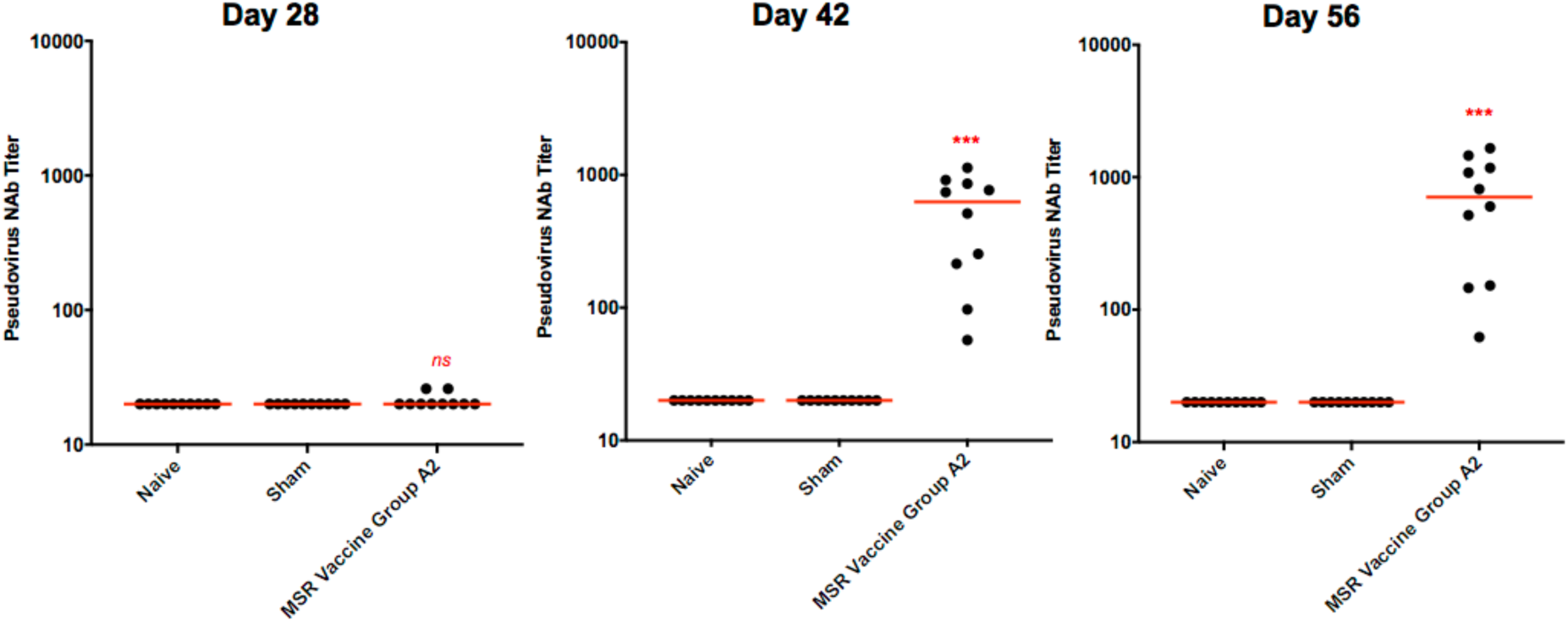
MSR add-in-time vaccines induced neutralizing antibodies (Nab) against a SARS-CoV-2 pseudovirus. Functional antibodies from sera of vaccinated mice were assessed by *in vitro* pseudovirus neutrailzation assay for Group A2 (MSR, S1/S2/N, GM-CSF, MPLA) add-in-timeand sham (MSR, GM-CSF, MPLA) vaccines on day 28, 42, and 56. Naïve mice were unvaccinated controls. Samples were plotted as median values (n=10).*, **, and*** represents p<0.05, p<0.01, and p<0.001, respectively.

## DISCUSSION

The global COVID-19 pandemic has accelerated research aiming to discover safe and effective vaccines against this pathogen, which likely will be needed to end the pandemic (*24*). Several vaccines are already in clinical trials, but multiple vaccines will likely be needed, and ones that can be easily and rapidly manufactured may be particularly useful (*25*). To address this need, we leveraged our biomaterial vaccine technology that is based on mesoporous silica. Synthetic amorphous silica is known to have good biocompatibility, supporting its development as a versatile platform for clinical applications. Further, the GM-CSF and MPLA used in the MSR vaccine have a track record of safe use in humans. Local release of these agents from a biomaterial vaccine allows small quantities of these agents to be used (e.g., 1 μg of GM-CSF per vaccine), minimizing concerns regarding systemic side-effects or toxicity. The sustained presentation of antigen and adjuvant made possible by the MSR vaccine, along with the accumulation of large numbers of dendritic cells at the vaccine site, has been previously demonstrated to yield robust and durable immune response in the context of oncology, as well as infectious diseases (*18-21, 26*). One of the great benefits of this technology is that the modularity enables plug-and-play versatility to quickly incorporate any antigen to manufacture and scale easily. Here we utilized recombinant protein subunit vaccines, as they allow ready control over epitopes and amount of antigen delivered, and the potential to fine-tune immune responses with adjuvants.

The results of these studies demonstrate that adding SARS-CoV-2 antigens to shell vaccines prior to immunization (add-in-time) induces robust humoral response, similar over time as those obtained with pre-manufactured lyophilized MSR vaccines. This add-in-time approach allows one to pre-manufacture shell vaccines (no antigen) in high quantities, store, and simply add antigen when situations such as this pandemic occur, and one requires a rapidly deployed vaccine. The kinetics of the IgG response were slightly slower with the add-in-time version of the MSR vaccine, and this could be a result of an insufficient times allowed for antigen adsorption prior to vaccination. Similar humoral responses were found with the two antigen doses tested in this study, suggesting the lower (1 μg) dose was already saturating.

Strong neutralizing responses were found in the sera of animals vaccinated with the various MSR vaccines, as measured by a SARS-CoV-2 pseudovirus assay. The Group A1 and A2 vaccines showed weaker activity before the boost on day 28, but strong activity was found on days 42 and 56. In contrast, strong neutralization activity was found on day 28 for the Group B, C, and D vaccines. Since the differentiating factor between these vaccines and Groups A1 and A2 was the addition of RBD in place of the N protein, we hypothesize this led to more effective neutralization. This result supports the relevance of the RBD in SARS-CoV-2 vaccines. Analysis of sera obtained from vaccinated animals at later time points will be tested in the future to provide a more in-depth analysis.

In summary, the data overall demonstrate the potential of the MSR vaccine to rapidly generate potent and protective humoral responses to SARS-CoV-2.

## ACKNOWLEDGEMENTS

We thank M. O. Dellacherie and M. Karkada for generous advice.

## Author contributions

F.L. and M.J.C. designed the experiments which were directed by E.J.D., D.H.B., and D.J.M. Vaccine manufacture and *in vitro* experiments were performed and analyzed by F.L., B.T.S., J.Y., M.J.C., C.Y., and D.W. *In vivo* work was performed by F.L. The manuscript was written by F.L. and B.T.S. All authors critically reviewed.

## REFERENCES

1. F. Wu, S. Zhao, B. Yu, Y. M. Chen, W. Wang, Z. G. Song, Y. Hu, Z. W. Tao, J. H. Tian, Y. Y. Pei, M. L. Yuan, Y. L. Zhang, F. H. Dai, Y. Liu, Q. M. Wang, J. J. Zheng, L. Xu, E. C. Holmes, Y. Z. Zhang, A new coronavirus associated with human respiratory disease in China. Nature 579, 265–269 (2020).

2. Q. Li, X. Guan, P. Wu, X. Wang, L. Zhou, Y. Tong, R. Ren, K. S. M. Leung, E. H. Y. Lau, J. Y. Wong, X. Xing, N. Xiang, Y. Wu, C. Li, Q. Chen, D. Li, T. Liu, J. Zhao, M. Liu, W. Tu, C. Chen, L. Jin, R. Yang, Q. Wang, S. Zhou, R. Wang, H. Liu, Y. Luo, Y. Liu, G. Shao, H. Li, Z. Tao, Y. Yang, Z. Deng, B. Liu, Z. Ma, Y. Zhang, G. Shi, T. T. Y. Lam, J. T. Wu, G. F. Gao, B. J. Cowling, B. Yang, G. M. Leung, Z. Feng, Early Transmission Dynamics in Wuhan, China, of Novel Coronavirus-Infected Pneumonia. N Engl J Med 382, 1199–1207 (2020).

3. N. Chen, M. Zhou, X. Dong, J. Qu, F. Gong, Y. Han, Y. Qiu, J. Wang, Y. Liu, Y. Wei, J. Xia, T. Yu, X. Zhang, L. Zhang, Epidemiological and clinical characteristics of 99 cases of 2019 novel coronavirus pneumonia in Wuhan, China: a descriptive study. Lancet 395, 507–513 (2020).

4. N. Zhu, D. Zhang, W. Wang, X. Li, B. Yang, J. Song, X. Zhao, B. Huang, W. Shi, R. Lu, P. Niu, F. Zhan, X. Ma, D. Wang, W. Xu, G. Wu, G. F. Gao, W. Tan, I. China Novel Coronavirus, T. Research, A Novel Coronavirus from Patients with Pneumonia in China, 2019. N Engl J Med 382, 727–733 (2020).

5. World Health Organization. WHO Coronavirus Disease (COVID-19) Dashboard. https://covid19.who.int/?gclid=EAIaIQobChMI5amijb_y6QIViovICh2yagafEAAYASABEgIv8PD_BwE (2020).

6. L. Ferretti, C. Wymant, M. Kendall, L. Zhao, A. Nurtay, L. Abeler-Dorner, M. Parker, D. Bonsall, C. Fraser, Quantifying SARS-CoV-2 transmission suggests epidemic control with digital contact tracing. Science 368, (2020).

7. A. Casadevall, M. J. Joyner, L. A. Pirofski, A Randomized Trial of Convalescent Plasma for COVID-19-Potentially Hopeful Signals. JAMA, (2020).

8. Y. Zhang, Q. Xu, Z. Sun, L. Zhou, Current targeted therapeutics against COVID-19: Based on first-line experience in China. Pharmacol Res 157, 104854 (2020).

9. F. Amanat, F. Krammer, SARS-CoV-2 Vaccines: Status Report. Immunity 52, 583–589 (2020).

10. K. Dhama, K. Sharun, R. Tiwari, M. Dadar, Y. S. Malik, K. P. Singh, W. Chaicumpa, COVID-19, an emerging coronavirus infection: advances and prospects in designing and developing vaccines, immunotherapeutics, and therapeutics. Hum Vaccin Immunother, 1–7 (2020).

11. W. Tai, L. He, X. Zhang, J. Pu, D. Voronin, S. Jiang, Y. Zhou, L. Du, Characterization of the receptor-binding domain (RBD) of 2019 novel coronavirus: implication for development of RBD protein as a viral attachment inhibitor and vaccine. Cell Mol Immunol 17, 613–620 (2020).

12. H. Bisht, A. Roberts, L. Vogel, K. Subbarao, B. Moss, Neutralizing antibody and protective immunity to SARS coronavirus infection of mice induced by a soluble recombinant polypeptide containing an N-terminal segment of the spike glycoprotein. Virology 334, 160–165 (2005).

13. J. D. Berry, K. Hay, J. M. Rini, M. Yu, L. Wang, F. A. Plummer, C. R. Corbett, A. Andonov, Neutralizing epitopes of the SARS-CoV S-protein cluster independent of repertoire, antigen structure or mAb technology. MAbs 2, 53–66 (2010).

14. L. Du, G. Zhao, Y. He, Y. Guo, B. J. Zheng, S. Jiang, Y. Zhou, Receptor-binding domain of SARS-CoV spike protein induces long-term protective immunity in an animal model. Vaccine 25, 2832–2838 (2007).

15. L. Du, Y. He, Y. Zhou, S. Liu, B. J. Zheng, S. Jiang, The spike protein of SARS-CoV--a target for vaccine and therapeutic development. Nat Rev Microbiol 7, 226–236 (2009).

16. J. Shang, Y. Wan, C. Luo, G. Ye, Q. Geng, A. Auerbach, F. Li, Cell entry mechanisms of SARS-CoV-2. Proc Natl Acad Sci U S A 117, 11727–11734 (2020).

17. X. Ou, Y. Liu, X. Lei, P. Li, D. Mi, L. Ren, L. Guo, R. Guo, T. Chen, J. Hu, Z. Xiang, Z. Mu, X. Chen, J. Chen, K. Hu, Q. Jin, J. Wang, Z. Qian, Characterization of spike glycoprotein of SARS-CoV-2 on virus entry and its immune cross-reactivity with SARS-CoV. Nat Commun 11, 1620 (2020).

18. J. Kim, W. A. Li, Y. Choi, S. A. Lewin, C. S. Verbeke, G. Dranoff, D. J. Mooney, Injectable, spontaneously assembling, inorganic scaffolds modulate immune cells in vivo and increase vaccine efficacy. Nat Biotechnol 33, 64–72 (2015).

19. M.O. Dellacherie et al., https://www.biorxiv.org/content/10.1101/2020.03.17.993808v1 (2020).

20. O. A. Ali, D. Emerich, G. Dranoff, D. J. Mooney, In situ regulation of DC subsets and T cells mediates tumor regression in mice. Sci Transl Med 1, 8ra19 (2009).

21. M. Super et al., https://www.biorxiv.org/content/10.1101/2020.02.25.964601v1 (2020).

22. C. R. Casella, T. C. Mitchell, Putting endotoxin to work for us: monophosphoryl lipid A as a safe and effective vaccine adjuvant. Cell Mol Life Sci 65, 3231–3240 (2008).

23. Z. Y. Yang, W. P. Kong, Y. Huang, A. Roberts, B. R. Murphy, K. Subbarao, G. J. Nabel, A DNA vaccine induces SARS coronavirus neutralization and protective immunity in mice. Nature 428, 561–564 (2004).

24. N. Lurie, M. Saville, R. Hatchett, J. Halton, Developing Covid-19 vaccines at pandemic speed. N Engl J Med 382, 1969–1973 (2020).

25. L. Corey, J. R. Mascola, A. S. Fauci, F. S. Collins, A strategic approach to COVID-19 vaccine R&D. Science 368, 948–950 (2020).

26. A. W. Li, M. C. Sobral, S. Badrinath, Y. Choi, A. Graveline, A. G. Stafford, J. C. Weaver, M. O. Dellacherie, T. Y. Shih, O. A. Ali, J. Kim, K. W. Wucherpfennig, D. J. Mooney, A facile approach to enhance antigen response for personalized cancer vaccination. Nat Mater 17, 528–534 (2018).

